# *Enterococcus faecalis and pathogenic streptococci* inactivate daptomycin by releasing phospholipids

**DOI:** 10.1101/149971

**Authors:** Elizabeth V. K. Ledger, Vera Pader, Andrew M. Edwards

**Affiliations:** MRC Centre for Molecular Bacteriology and Infection, Imperial College London, Armstrong Rd, London, SW7 2AZ, UK.

## Abstract

Daptomycin is a lipopeptide antibiotic with activity against Gram-positive bacteria. We have shown previously that *Staphylococcus aureus* can survive daptomycin exposure by releasing membrane phospholipids that inactivate the antibiotic. To determine whether other pathogens possess this defence mechanism, phospholipid release and daptomycin activity were measured after incubation of *Staphylococcus epidermidis*, Group A or B streptococci, *Streptococcus gordonii* or *Enterococcus faecalis* with the antibiotic. All bacteria released phospholipid in response to daptomycin, which resulted in at least partial inactivation of the antibiotic. However, *E. faecalis* showed the highest levels of lipid release and daptomycin inactivation. As shown previously for *S. aureus*, phospholipid release by *E. faecalis* was inhibited by the lipid biosynthesis inhibitor platensimycin. In conclusion, several pathogenic Gram-positive bacteria, including *E. faecalis*, inactivate daptomycin by releasing phospholipids, which may contribute to the failure of daptomycin to resolve infections caused by these pathogens.

## Manuscript text

Daptomycin is a lipopeptide antibiotic used as a last resort in the treatment of infections caused by methicillin-resistant *Staphylococcus aureus* (MRSA) and vancomycin-resistant enterococci (VRE) [1-3]. The use of daptomycin is becoming more common, with prescriptions increasing 72 % between 2012 and 2015 in the UK [4]. Daptomycin is the only lipopeptide antibiotic used clinically and functions in a similar manner to antimicrobial peptides [5]. The antibiotic inserts into the membrane of Gram-positive bacteria by targeting phosphatidylglycerol, where it forms oligomeric complexes [6-8]. The precise mechanism by which the antibiotic kills bacteria is unclear, but involves depolarisation of the bacterial membrane and inhibition of cell wall biosynthesis without causing lysis [8-13]. Although daptomycin resistance is rare, treatment failure occurs in up to 30 % of staphylococcal infections and 23 % of enterococcal infections [14,15]. Failure rates are highest in invasive infections such as bacteraemia or osteomyelitis, with rates of 24 % and 33 % respectively, resulting in poor patient prognoses [14]. Understanding the reasons for this treatment failure is crucial to improving the effectiveness of daptomycin treatment.

We recently discovered that *S. aureus* has a transient defence mechanism against daptomycin, which contributed to treatment failure in a murine model of invasive infection [16]. In response to the antibiotic, phospholipids were released from the cell membrane which sequestered daptomycin and abrogated its bactericidal activity [16]. Phospholipid release occurred via an active process, which was blocked by the lipid biosynthesis inhibitor platensimycin [16,17]. In addition to daptomycin, phospholipid shedding also provided protection against the antimicrobial peptides nisin and melittin, suggesting a general defence against membrane-targeting antimicrobials [16].

It is currently unknown whether other Gram-positive bacteria release phospholipids in response to daptomycin, although membrane vesicles have been observed on the surface of *E. faecalis* cells exposed to daptomycin [18]. In addition, there is growing evidence that other Gram-positive pathogens, including Group A streptococci (GAS) and group B streptococci (GBS), release phospholipids from their surfaces in the form of extracellular vesicles [19,20]. Production of these membrane vesicles is increased in the presence of antimicrobials and, at least for GAS, are rich in phosphatidylglycerol, which was shown to be essential for daptomycin inactivation by *S. aureus* [16,19,21]. Therefore, we hypothesised that phospholipid release is a common strategy amongst Gram-positive pathogens to resist membrane-acting antimicrobials.

Given the increasing use of daptomycin to treat enterococcal infections, the primary aim of this work was to determine whether enterococci release membrane phospholipids that inactivate the antibiotic. We also examined pathogenic streptococci, and *S. epidermidis*, as the rising tide of antibiotic resistance may necessitate the use of daptomycin to tackle these bacteria in the future.

We initially determined the daptomycin minimum inhibitory concentration (MIC) for a representative panel of Gram-positive pathogens: *S. aureus* SH1000 [22], *S. epidermidis* ATCC 12228 [23], GAS strain A40 [24]; GBS strains 515 [25] and COH1 [26]; *S. gordonii* strain Challis [27]; and *E. faecalis* strains JH2-2 [28] and OG1X [29]. Bacteria were grown in Muller Hinton Broth containing calcium (0.5 mM) and MIC determined by the broth microdilution approach [30]. The most susceptible species were the pathogenic GAS strain A40 (0.125 μg ml^-1^), and GBS strains 515 (0.5 μg ml^-1^) and COH1 (0.5 μg ml^-1^), whilst *S. aureus* (1 μg ml^-1^), *S. epidermidis* (1 μg ml^-1^), *S. gordonii* Challis (2-4 μg ml^-1^) *E. faecalis* strains OG1X (2 μg ml^-1^) and JH2-2 (4 μg ml^-1^) were the least susceptible.

To determine whether *E. faecalis* or streptococci respond to daptomycin by releasing membrane phospholipids, we exposed streptococci and enterococci (10^8^ CFU ml^-1^) to various supra-MIC concentrations of the antibiotic (5-40 μg ml^-1^) in Brain-Heart Infusion (0.5 mM CaCl_2_) broth at 37 °C under static conditions with 5% CO_2_ and measured bacterial survival, antibiotic activity and phospholipid release (Fig. 1c-h). Staphylococci were also exposed to daptomycin (5-40 μg ml^-1^), but in tryptic soy broth (TSB) containing 0.5 mM CaCl2 at 37 °C with shaking (180 RPM) (Fig. 1a,b).

**Fig. 1.**
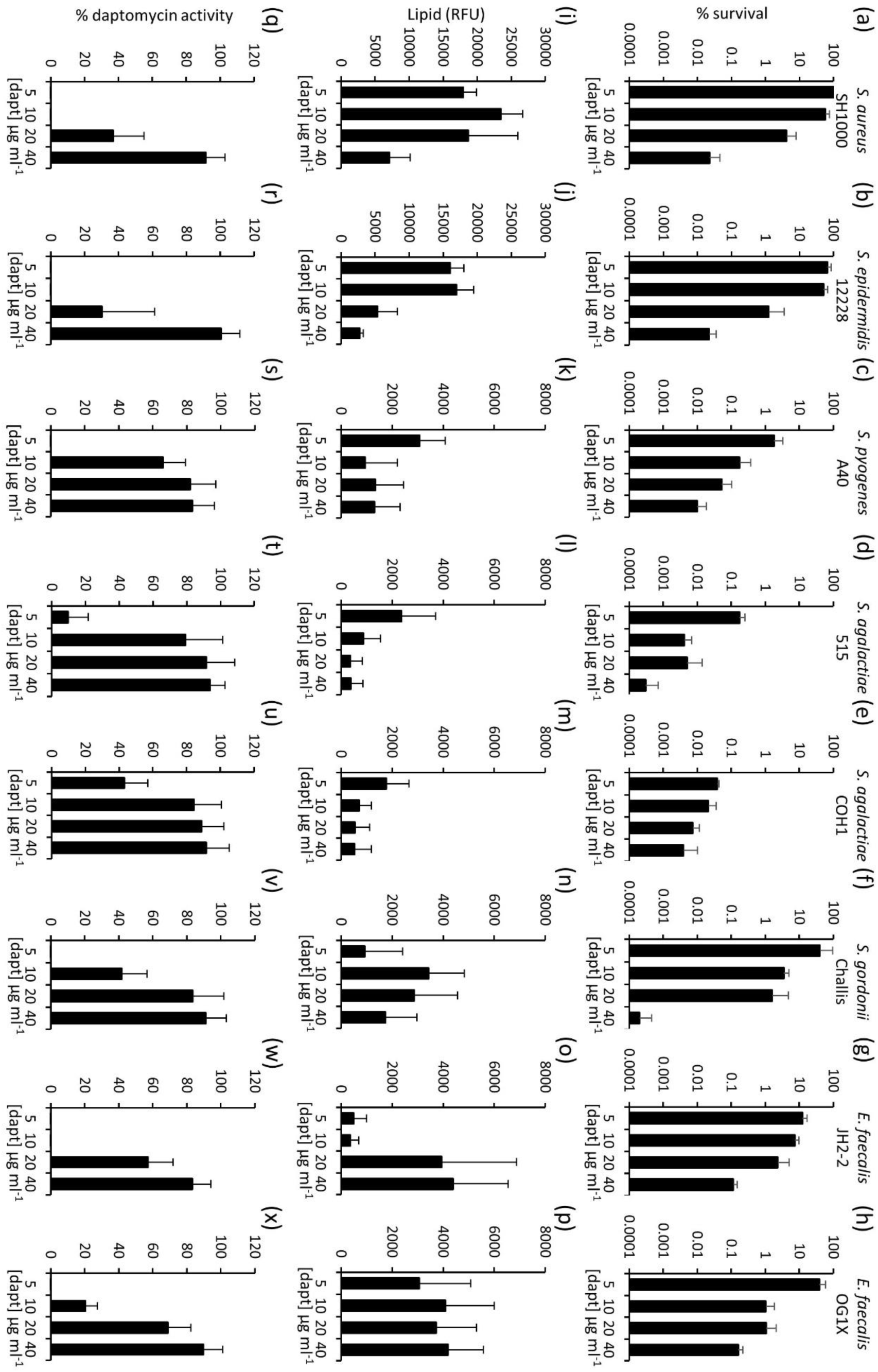
Streptococci and enterococci release phospholipid and inactivate daptomycin. (a-h), percentage survival of bacteria after 8 h incubation in broth containing the indicated concentrations of daptomycin. (i-p) the concentration of phospholipid in culture supernatants of bacteria exposed to daptomycin as determined by reactivity with a fluorescent dye (RFU: relative fluorescence units). Note the different Y-axis scale for staphylococci vs other bacteria. (q-x) relative percentage of daptomycin activity remaining in culture supernatants of bacteria exposed to daptomycin for 8 h. The activity of daptomycin incubated in culture medium only for 8 h was taken to be 100 %. For all data, the mean of 4 independent experiments are shown, and error bars represent the standard deviation of the mean.

For all strains, there was a dose-dependent decrease in survival after 8 h exposure to daptomycin, as assessed by CFU counts (Fig. 1a-h). However, as expected from the MIC data, survival of the two enterococcal strains, the staphylococci and *S. gordonii* was greater than survival of GAS or GBS strains at every concentration of daptomycin examined (Fig. 1a-h).

Using the phospholipid-reactive fluorescent dye FM-4-64 (Life Technologies), we found that daptomycin triggered phospholipid release from all of the bacteria examined, albeit to differing levels. The quantity of phospholipid released was much greater for staphylococci than the other species examined (Fig. 1i-p). However, for both staphylococci and streptococci, the quantity of phosphlipid released was lowest when the daptomycin concentration was highest, suggesting that the antibiotic may have killed the bacteria before they could release the lipid (Fig. 1i-p). By contrast, the enterococci released high levels of phospholipid in the presence of the highest concentrations of daptomycin (Fig. 1o,p). This may indicate different daptomycin concentration thresholds for triggering of phospholipid release.

To determine whether phospholipid release resulted in the inactivation of daptomycin, the activity of the antibiotic in the culture supernatants was measured using a previously described zone of inhibition assay [16] (Fig. 1q-x). Daptomycin was inactivated to varying degrees by the bacteria, depending on the concentration of the antibiotic used. However, both staphylococcal strains, both enterococcal strains, *S. gordonii* and the GAS strain completely inactivated daptomycin at 5 µg ml^-1^, but GBS strains only partially inactivated the antibiotic at this concentration. At 10 µg ml^-1^ daptomycin, only the staphylococci, *S. gordonii* and the enterococci showed significant inactivation of the antibiotic and at a concentration of 20 µg ml^-1^ daptomycin, only staphylococci and enterococci inactivated the antibiotic to any significant degree, with a loss of 30-60% of antibiotic activity. However, despite triggering phospholipid release, at 40 µg ml^-1^ daptomycin there was relatively little (<20%) inactivation of the antibiotic by any of the bacteria tested. Therefore, phospholipid release is finite and can be overcome with a sufficiently high dose of daptomycin.

These data extend our previous finding that *S. aureus* releases phospholipid in response to daptomycin and that this results in inactivation of the antibiotic by revealing a very similar phenotype for *S. epidermidis*. These data also support the previous observation that *E. faecalis* releases phospholipid in response to daptomycin [18], and show that this phospholipid release correlates with daptomycin inactivation and bacterial survival. Streptococci, particularly *S. gordonii*, also released phospholipid and inactivated daptomycin, albeit less efficiently than *E. faecalis*. Therefore, daptomycin-induced phospholipid release appears to be a conserved mechanism across Gram-positive pathogens.

Next, we wanted to explore whether the mechanism of phospholipid release and daptomycin inactivation by enterococci and streptococci was similar to that of *S. aureus*. Therefore, we undertook further experiments with *E. faecalis*, which was most efficient of the enterococci and streptococci at releasing phospholipid and inactivating daptomycin, and *S. aureus*, in which daptomycin-triggered phospholipid release has been well characterised [16].

In *S. aureus*, daptomycin-triggered phospholipid release is an active process that requires energy, as well as protein and lipid biosynthesis [16]. To determine whether phospholipid release by *E. faecalis* exposed to daptomycin was occurring via an active process, or simply a consequence of damage caused by the antibiotic, bacteria were exposed to the antibiotic in the presence or absence of a sub-inhibitory concentration of the phospholipid biosynthesis inhibitor platensimycin [17]. As described previously, exposure of *S. aureus* to daptomycin (10 µg ml^-1^) resulted in increased phospholipid in the supernatant but this was significantly reduced in the presence of platensimycin at half the MIC (0.25 μg ml^-1^) (Fig. 2a). Similarly, phospholipid was released upon exposure of *E. faecalis* to daptomycin (10 µg ml^-1^), but this was blocked when platensimycin was present at half the MIC (0.5 μg ml^-1^) (Fig. 2b). The presence of platensimycin prevented *S. aureus* from inactivating daptomycin (Fig. 2c) and significantly reduced the ability of *E. faecalis* to inactivate daptomycin (Fig. 2d). This confirmed that daptomycin-induced phospholipid release by *E. faecalis* is an active process that requires *de novo* lipid biosynthesis and is not simply a consequence of membrane damage caused by the antibiotic. The ability of platensimycin to block phospholipid release and prevent daptomycin inactivation by *E. faecalis* also provided strong evidence that, as for *S. aureus*, daptomycin activity is blocked by the phospholipid in the supernatant. However, it was necessary to rule out an alternative hypothesis; that the loss daptomycin inactivation was simply due to binding of the antibiotic to the bacterial surface.

**Fig. 2.**
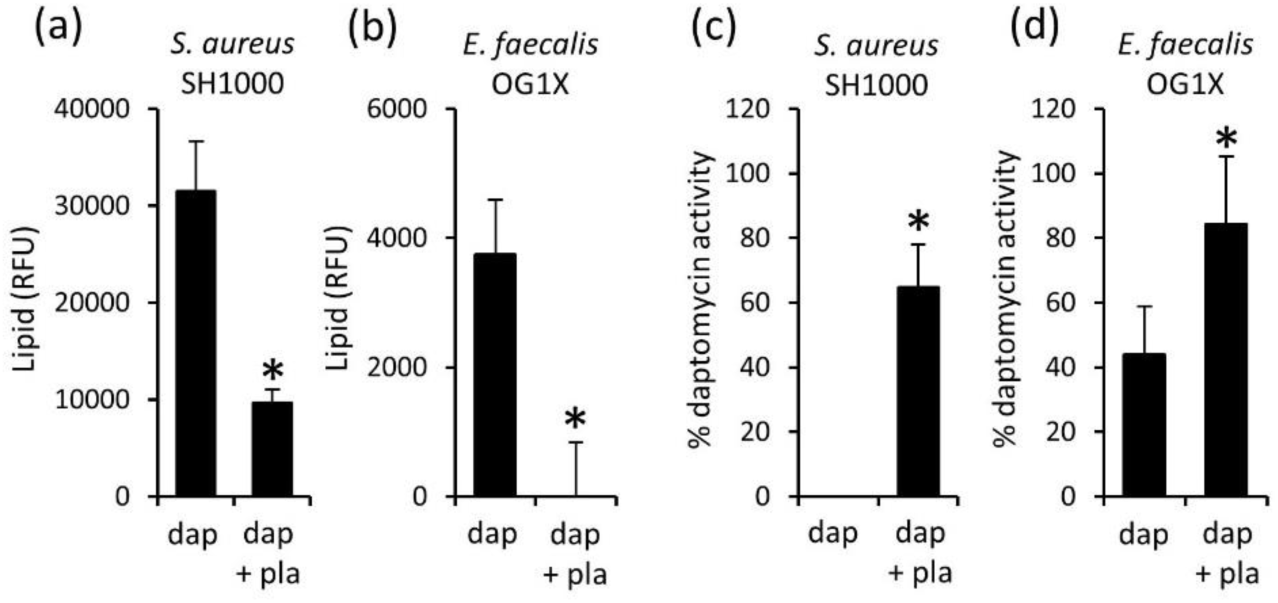
*De novo* lipid biosynthesis is required for enterococcal inactivation of daptomycin. Phospholipid concentration (RFU) in culture supernatants from *S. aureus* (a) or E. faecalis OG1X (b) incubated for 8 h in media containing daptomycin (10 µg ml^-1^) only (dap) or both daptomycin and 0.5 X MIC platensimicin (dap + pla). (c,d) relative % daptomycin activity in supernatants from cultures described in (a) and (b), respectively. Data in (a) and (b) were analysed using a one-way ANOVA with Tukey’s post-hoc test. Data in (c) and (d) were analysed by a Student’s *t*-test. *P=<0.05.

To measure binding of daptomycin to bacteria, daptomycin was labelled with the Bodipy fluorophore (Life Technologies) as described previously [11,16]. As reported previously, a killing assay with *E. facealis* indicated that the labelled antibiotic had slightly altered bactericidal activity relative to unlabelled daptomycin [11] (Fig 3a). However, as described above for unlabelled antibiotic (Fig. 1q,x), the activity of the antibiotic decreased after incubation with *E. faecalis* or *S. aureus* (Fig. 3b), confirming that the Bodipy label does not significantly affect the interaction of the antibiotic with the bacteria studied.

**Fig. 3.**
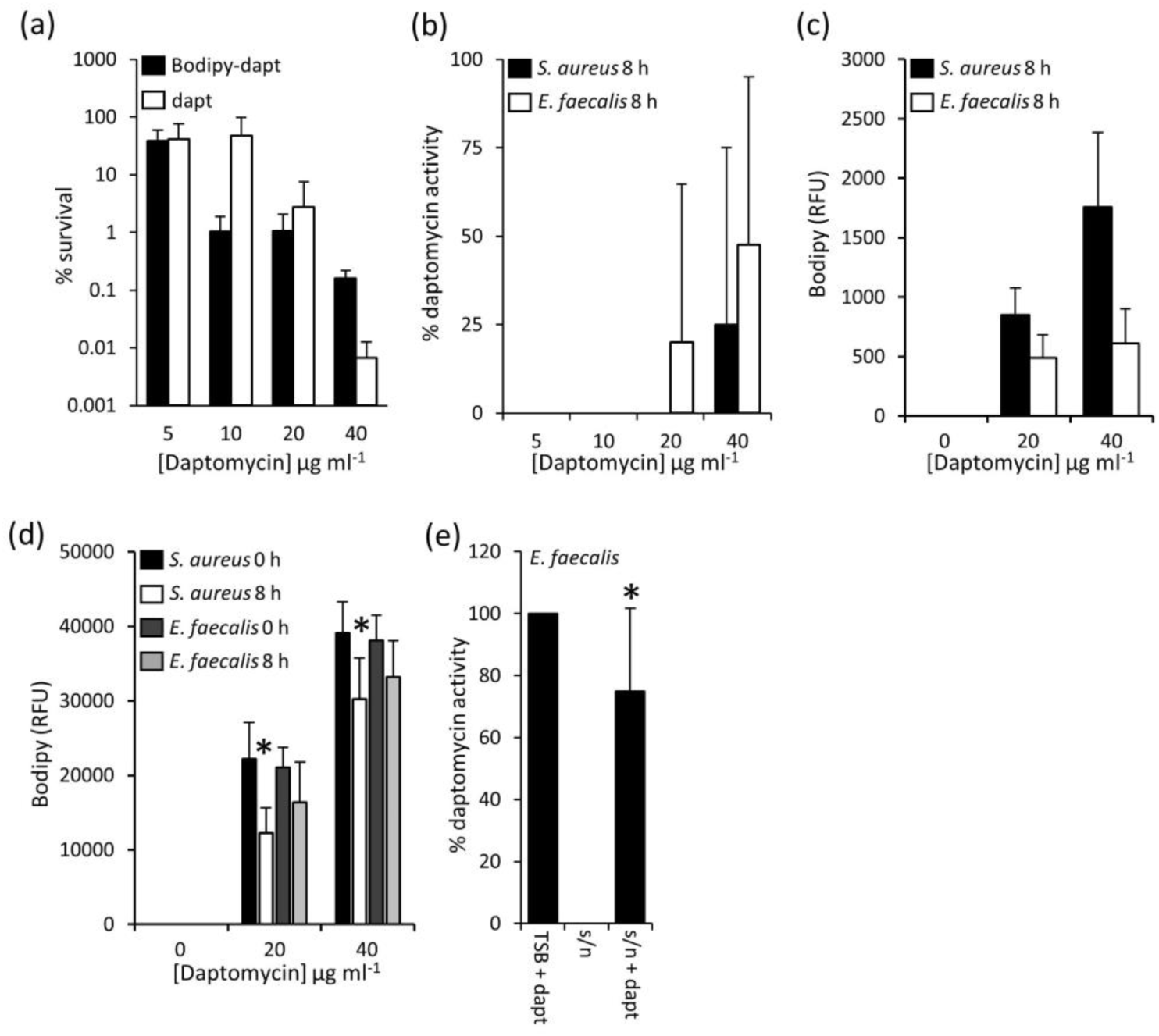
Loss of daptomycin activity in supernatant is not due to antibiotic binding to bacteria. (a) percentage survival of *E. faecalis* OG1X incubated with various concentrations of daptomycin (dapt) or Bodipy-daptomycin (Bodipy-dapt) for 8 h. (b) relative percentage daptomycin activity in culture supernatants described in (a). (c) binding of Bodipy-daptomycin to *S. aureus* or *E. faecalis* OG1X after 8 h incubation in media containing the indicated concentration of the labelled antibiotic. (d) quantification of Bodipy-daptomycin (RFU) remaining in culture supernatants from *S. aureus* or *E. faecalis* OG1X, after 8 h incubation with Bodipy-daptomycin as described in (c) * indicates significantly different from 0 h time point. (e) relative percentage activity of daptomycin (5 μg ml^-1^) activity in TSB only (TSB + dapt), in the supernatant from *E. faecalis* incubated with daptomycin for 8 h (s/n) and after the addition of 5 μg ml^-1^ daptomycin to the supernatant from *E. faecalis* incubated with daptomycin for 8 h (s/n + dapt). Data in (d) and (e) were analysed by a two-way ANOVA with Tukey’s post-hoc test. Graphs show the mean average and, where shown, error bars represent the standard deviation of the mean. For each panel *P=<0.05.

After 8 h incubation with Bodipy-daptomycin, bacterial cells were pelleted and the fluorescence of both the cells and the supernatants was measured separately using a Tecan microplate reader with excitation at 502 nm and emission at 510 nm. Antibiotic attachment to the *E. faecalis* cellular fraction was similar for both Bodipy-daptomycin concentrations examined, suggesting saturated binding to cells (Fig. 3c). However, most of the antibiotic remained in the supernatant (Fig. 3d). By comparison, Bodipy-daptomycin bound *S. aureus* more strongly than *E. faecalis*, with higher levels of fluorescence associated with bacterial cells and a corresponding drop in the fluorescence of the supernatant (Fig. 3c,d). This difference in antibiotic binding may explain why the daptomycin MIC of the *E. faecalis* strains used here (2-4 µg ml^-1^) is higher than that of the *S. aureus* strain examined (1 µg ml^-1^) and why daptomycin triggers greater phospholipid release from staphylococci than enterococci.

Together, these data confirmed that the loss of daptomycin activity in *E. faecalis* cultures was not due to binding of the antibiotic to the bacterial surface or the plastic vessels used in the assays. However, as a final confirmation that phospholipid released from *E. faecalis* inactivated daptomycin, we exposed the bacterium to daptomycin (5 µg ml^-1^) to trigger phospholipid release, collected the cell-free culture supernatant and added a second dose of the antibiotic (5 µg ml^-1^). The culture supernatant containing the released phospholipids significantly reduced the activity of the second dose of daptomycin (by ^~^25%, Fig. 3e). Therefore, as described for *S. aureus*, the release of phospholipids by *E. faecalis* in response to daptomycin inactivates the antibiotic. The data described above also indicate that several species of streptococci release phospholipids in response to daptomycin, which inactivate the antibiotic, albeit to a lesser extent than *E. faecalis* or *S. aureus.*

Streptococci and enterococci cause a range of serious diseases, including septicaemia and endocarditis, which can be treated by daptomycin, especially when the pathogen is multi-drug resistant or the patient has a β-lactam allergy [1,31]. The presence of this defence mechanism in a variety of clinically-relevant Gram-positive bacteria indicates that it is conserved and could be a viable target to improve the effectiveness of daptomycin therapy against these pathogens.

In this work, we focussed on daptomycin because it is a last resort antibiotic and is associated with high rates of treatment failure. However, whilst daptomycin use is increasing, it is very unlikely to have provided the selection pressure for the evolution of the phospholipid release defence mechanism described here and previously [16]. Since cationic antimicrobial peptides (cAMPs) act via a similar mechanism to daptomycin in targeting the Gram-positive cell membrane [5] we hypothesise that these host defence molecules have likely driven the evolution of phospholipid release as a defence mechanism.

The discovery of phospholipid release in several Gram-positive pathogens expands our growing appreciation of broad-spectrum extracellular defence mechanisms that protect bacteria against antibiotics or host defences. For example, previous work has shown that the production of outer-membrane vesicles by *E. coli* can protect against membrane-acting antimicrobials such as polymixin E and colistin [32], whilst another report revealed that lipochalins released by *Burkholderia* can sequester several different antibiotics [33]. These findings underline the complex nature of innate antibiotic resistance, but also provide opportunities for mechanistic insight and improved therapeutic approaches. For example, in this report and previously, we have shown that inhibition of phospholipid biosynthesis using platensimycin prevents the inactivation of daptomycin by both *S. aureus* and *E. faecalis* [16]. Although platensimycin has not entered clinical trials due to poor pharmacokinetic properties [17,34], other inhibitors of lipid biosynthesis are in clinical development [35]. Therefore, the use of daptomycin in combination with lipid biosynthesis inhibitors may provide an effective way of enhancing treatment outcomes compared to the lipopeptide antibiotic alone.

In summary, we have demonstrated that *Enterococcus faecalis* releases phospholipids in response to daptomycin via an active mechanism requiring *de novo* lipid biosynthesis and that these phospholipids inactivate daptomycin. Pathogenic streptococci also appear to be capable of inactivating daptomycin by releasing phospholipids, indicating that this mechanism is conserved amongst Gram-positive pathogens.

## Acknowledgements

Mal Horsburgh (University of Liverpool) and Angela Nobbs (University of Bristol) are acknowledged for kindly proving strains. E.V.K.L. is supported by a Wellcome Trust PhD studentship through an award to Imperial College. A.M.E. acknowledges funding from the Royal Society, Department of Medicine and from the Imperial NIHR Biomedical Research Centre, Imperial College London.

